# Mechanistic basis for decreased antimicrobial susceptibility in a clinical isolate of *Neisseria gonorrhoeae* possessing a mosaic-like *mtr* efflux pump locus

**DOI:** 10.1101/448712

**Authors:** Corinne E. Rouquette-Loughlin, Jennifer L. Reimche, Jacqueline T. Balthazar, Vijaya Dhulipala, Kim M. Gernert, Ellen N. Kersh, Cau D. Pham, Kevin Pettus, A. Jeanine Abrams, David L. Trees, Sancta St Cyr, William M. Shafer

## Abstract

Recent reports suggest that mosaic-like sequences within the *mtr* (*m*ultiple *t*ransferable *r*esistance) efflux pump locus of *Neisseria gonorrhoeae* likely originating from commensal *Neisseria sp.* by transformation can increase the ability of gonococci to resist structurally diverse antimicrobials. Thus, acquisition of numerous nucleotide changes within the *mtrR* gene encoding the transcriptional repressor (MtrR) of the *mtrCDE* efflux pump-encoding operon or overlapping promoter region for both along with those that cause amino acid changes in the MtrD transporter protein were recently reported to decrease gonococcal susceptibility to numerous antimicrobials, including azithromycin (Azi) (**Wadsworth *et al.* 2018. MBio. doi.org/10.1128/mBio.01419-18**). We performed detailed genetic and molecular studies to define the mechanistic basis for why such strains can exhibit decreased susceptibility to MtrCDE antimicrobial substrates including Azi. We report that a strong *cis*-acting transcriptional impact of a single nucleotide change within the -35 hexamer of the *mtrCDE* promoter as well gain-of-function amino acid changes at the *C*-terminal region of MtrD can mechanistically account for the decreased antimicrobial susceptibility of gonococci with a mosaic-like *mtr* locus.

**IMPORTANCE:** Historically, after introduction of an antibiotic for treatment of gonorrhea, strains of *N. gonorrhoeae* emerge that display clinical resistance due to spontaneous mutation or acquisition of resistance genes. Genetic exchange between members of the *Neisseria* genus occurring by transformation can cause significant changes in gonococci that impact the structure of an antibiotic target or expression of genes involved in resistance. The results presented herein provide a framework for understanding how mosaic-like DNA sequences from commensal *Neisseria* that recombine within the gonococcal *mtr* efflux pump locus function to decrease bacterial susceptibility to antimicrobials including antibiotics used in therapy of gonorrhea.

## INTRODUCTION

*Neisseria gonorrhoeae* is the etiologic agent of the sexually transmitted infection (STI) gonorrhea. Gonorrhea is the second most reported condition in the USA (468,514 cases were reported in 2016) (1) and a major worldwide public health problem given its estimated yearly incidence of 78 million of infections (2). Historically, the gonococcus has developed resistance to all drugs used for treatment since the introduction of sulfonamides in the late 1930s (3) and concern exists that without new effective antibiotics some gonorrheal infections in the future may be untreatable (4, 5). Currently, a dual antibiotic treatment regimen of ceftriaxone (Cro) (single intramuscular injection of 250-500 mg) and azithromycin (Azi) (single oral dose of 1-2 g dose) is used in many western countries (6, 7), but their continued efficacy for use in curing gonorrheal infections is threatened as strains resistant to either or both antibiotics have emerged in the past decade (8-11).

The gonococcus has adapted numerous strategies to survive attacks by antimicrobials, including the use of multidrug efflux pumps to export toxic compounds (3, 12, 13). Five gonococcal efflux pumps that export a wide range of substrates have been described (13). Of these, the best studied efflux pump is MtrCDE, which belongs to the resistance-nodulation-division family possessed by many Gram-negative bacteria. MtrCDE captures and exports structurally diverse, but generally amphipathic, antimicrobial agents including macrolides, beta-lactams, cationic antimicrobial peptides, dyes and detergents (13). The contribution of the MtrCDE efflux pump in antimicrobial resistance expressed by gonococci can be enhanced by *cis*- or *trans*-acting mutations that result in over-expression of the *mtrCDE* efflux pump operon (13). Importantly, over-production of the MtrCDE efflux pump due to relief of transcriptional repression of *mtrCDE* can contribute to clinically relevant levels of resistance to beta-lactams and macrolides (13).

Expression of *mtrCDE* in wild-type (WT) gonococci is subject to repression by MtrR (14, 15) and, in the presence of an inducer, activation by MtrA (16). Both MtrA and MtrR bind to regions within a 250 bp sequence that contains overlapping, divergent promoters for *mtrR* and *mtrCDE* transcription (17). Loss of MtrR repression of *mtrCDE* can result from point mutations in the MtrR-binding site (14), a single base pair (bp) deletion within a 13 bp inverted repeat sequence in the *mtrR* promoter (18), a point mutation that creates a new *mtrCDE* promoter (19) or missense/nonsense mutations in the *mtrR* gene (20-22). In addition to decreasing gonococcal susceptibility to antibiotics, these regulatory mutations can also enhance the fitness of gonococci during experimental infection of the lower genital tract of female mice (23), which supports the concept that the MtrCDE efflux pump is of importance during infection due to its ability to export host-derived antimicrobials such as cationic antimicrobial peptides and progesterone (24).

While single site regulatory mutations impacting *mtrCDE* expression have been extensively studied (14, 19-22, 29), increasing evidence suggests that entry and recombination of donor DNA from commensal *Neisseria spp.* into the *mtr* locus can result in multiple nucleotide changes that can decrease gonococcal susceptibility to antimicrobials including Azi and Cro. Thus, the presence of mosaic-like sequences within the *mtrR* region likely resulting from transformation by DNA from *N. lactamica* or *N. meningitidis* has been reported in worldwide-isolated gonococcal strains (25-28, 30). Recent work by Wadsworth *et al.* (30) showed that gonococci bearing diverse mosaic-like sequences within the *mtrR/mtrCDE* promoter region have elevated expression of *mtr*-associated genes and decreased susceptibility to Azi. Importantly, mosaic-like sequences within the *mtrD* inner membrane transporter protein-encoding gene showed strong linkage disequilibrium and epistatic effects that likely enhance the activity of the MtrCDE efflux pump (30). Taken together, the available information strongly suggest that mosaic-like sequences in the *mtr* locus can result in increased expression of the *mtrCDE* efflux pump operon as well as providing a gain-of-function property to MtrD that enhances its ability to export antimicrobials. In this study, we examined gonococcal clinical isolates that possess a mosaic-like *mtr* locus similar to other clinical isolates (30). We describe both a single nucleotide change in the overlapping *mtrR/mtrCDE* promoters and a likely mechanism for its impact on gene transcription, and amino acid changes in the C-terminal domain of MtrD that were linked to the decreased antimicrobial susceptibility phenotype expressed by gonococci with a mosaic-like *mtr* locus.

## RESULTS

### Importance of the MtrCDE efflux pump in reduced susceptibility to Azi and other antimicrobials in gonococcal clinical isolates bearing a mosaic-like *mtr* locus

Public health laboratories associated with the Gonococcal Isolate Surveillance Project (GISP) alert the Centers for Disease Control and Prevention (CDC) to *N. gonorrhoeae* isolates if the Azi minimal inhibitory concentration (MIC) is 2 µg/ml or greater. High-level MICs to Azi (≥ 256 µg/ml [3]) is typically due to mutations in the four 23S ribosomal RNA (*rRNA*) genes (31). However, in recent years it has become apparent that gonococci can display a so-called “less Azi-susceptible” phenotype characterized by MIC values of 2-4 µg/ml (3, 25-30) that does not involve 23S *rRNA* mutations. This less Azi-susceptible property may help gonococci escape the action of this macrolide during treatment, especially at extra-genital sites of infection (e.g., the pharyngeal mucosa) where the pharmacokinetic properties of antibiotics are not optimal (32). It is therefore important to define how gonococci can develop decreased Azi-susceptibility in the absence of 23S *rRNA* mutations as this could result in clinical failure of this macrolide during certain infections.

In order to study emergence of gonococcal clinical isolates with reduced susceptibility to Azi and to ascertain the contribution, if any, of the MtrCDE efflux pump system to this phenotype, we analyzed eight clinical strains collected in 2014 that expressed a less Azi-susceptible property (MIC of 2 µg/ml); details of these strains are provided in Materials and Methods and Table S1. Whole genome sequencing (WGS) performed on these eight strains and detailed bioinformatic analysis revealed that they lacked known 23S *rRNA* mutations associated with high levels of Azi resistance (data not presented). The sequence of the genes within the *mtr* locus of the 8 strains were identical, and they contained multiple nucleotide differences compared to antibiotic-sensitive reference strain FA19; the details of the WGS performed on these and other strains will be presented elsewhere (Soge and McLean, in preparation). Briefly, an alignment of the nucleotide sequence of the entire *mtr* locus in clinical strain LRRBGS0002 (hereafter termed CDC2), displaying a less Azi-susceptible phenotype with that of FA19 is shown in Fig. S1A. The nucleotide sequence of the *mtr* locus possessed by strain CDC2 was most dissimilar to FA19 (as well as three other gonococcal strains [MS11, FA1090 and H041]) in the *mtrR/mtrCDE* promoter region, the *mtrD* gene and the non-coding region between *mtrD* and *mtrE* (Fig. 1). A BLAST search against *Neisseria* (taxid: 482) nucleotide sequences in NCBI (blast.ncbi.nlm.nih.gov) determined that the entire *mtr* locus sequence possessed by CDC2 was most similar (95% identity) to that of *N. polysaccharea* M18661 (GenBank: CP031325.1). Analysis of a phylogenetic tree based on the *mtr* loci of CDC2, FA19, and three clinical strains (GCGS0276, GCGS0402, and GCGS0834) with mosaic-like *mtr* loci studied by Wadsworth *et al.* (30), indicated that at the *mtr* locus CDC2 was most similar to that possessed by strain GCGS0402 (Fig. S1B). In fact, the nucleotide sequence of the *mtr* locus of CDC2 and GCGS0402 was identical. Importantly, however, CDC2 lacked the Correia element (CE) (33) that is positioned adjacent to the *mtrR-mtrCDE* promoter region found in *mtr* mosaic-like strain GCGS0276 (30) (Fig. S1A).

**Figure 1.**
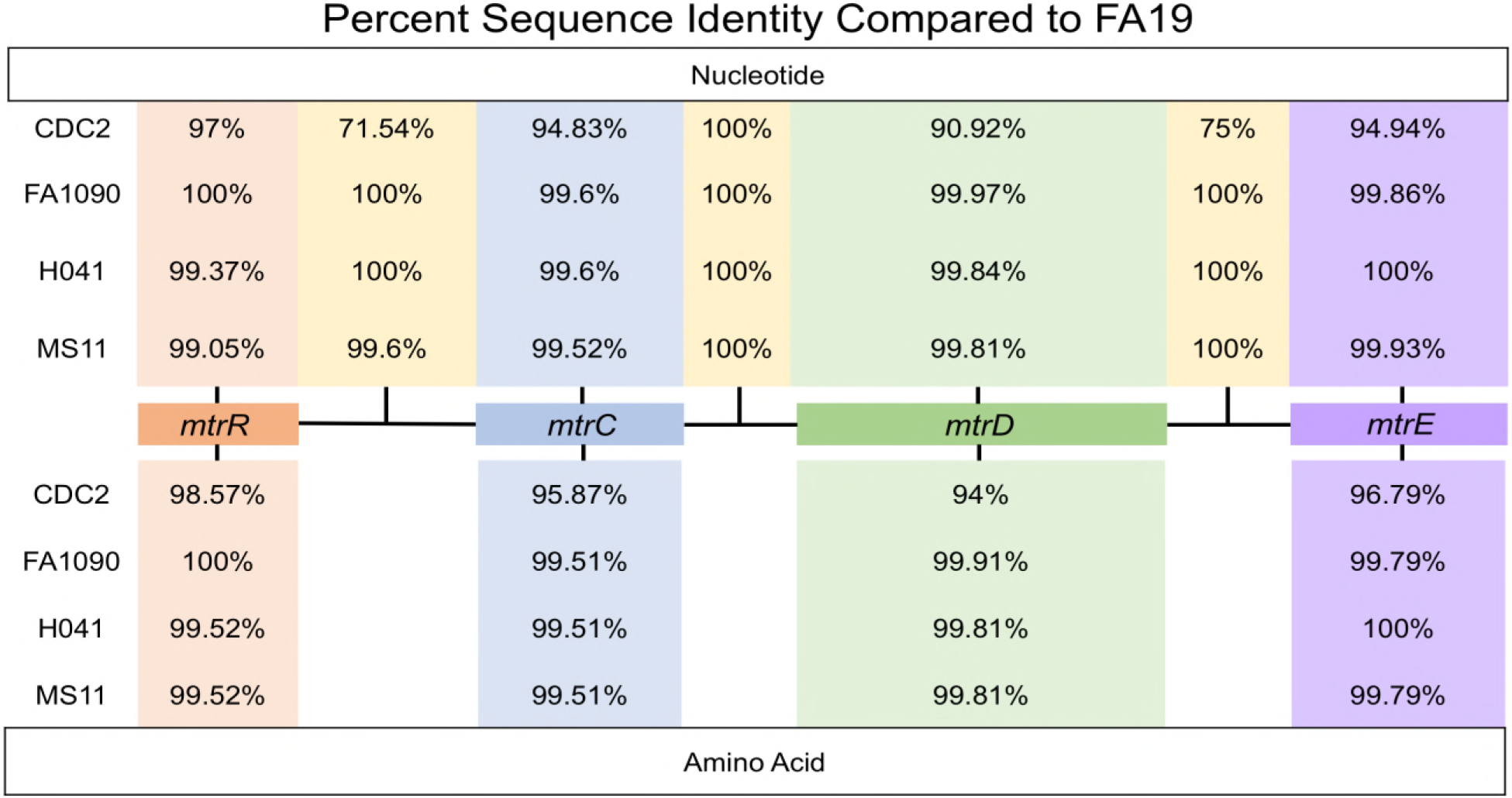
Shown are the nucleotide and amino acid sequence percent identities of genes and intervening regions of the *mtr* locus possessed by gonococcal strains with respect to FA19 (CP012026.1). Accession numbers are provided in Table S1. Clustal Omega multiple sequence alignments were performed using *N. gonorrhoeae* strains FA19, CDC2, FA1090, H041, and MS11. Alignments were generated for each *mtr* gene using their nucleotide and amino acid sequences, and nucleotide sequences were aligned for the intergenic regions. Pairwise identity matrices were calculated, and the pairwise identity values for each alignment are shown.

Since CDC2 contained numerous nucleotide sequence variations (with respect to FA19) in the *mtrR* coding region and the overlapping *mtrR/mtrCDE* promoter region, we hypothesized that it (as well as the seven other alert strains) might over-produce the MtrCDE pump leading to decreased susceptibility to Azi and other antimicrobials. In order to confirm that this efflux pump is required for antimicrobial resistance in CDC2, as it has been reported for non-*mtr* mosaic-like clinical isolates such as H041 (29), we created a mutant that lacked a functional MtrCDE efflux pump due to insertional inactivation (*mtrD::kan*) of the parental *mtrD* gene, which encodes the MtrD cytoplasmic membrane transporter. We also created other mutants that lacked functional MacAB or NorM efflux pumps to ascertain if their loss might also increase susceptibility of this strain to antimicrobials. Of these three efflux pumps, only loss of an active MtrCDE efflux pump rendered CDC2 and the other Azi alert isolates hyper-susceptible to Azi (Table S1). Similarly, for CDC2 (Table 1) and the other Azi alert strains (data not presented), only the loss of the MtrCDE efflux pump (see CR.99 in Tables S1 and S2) resulted in hyper-susceptibility to the tested antimicrobials.

**Table 1.**
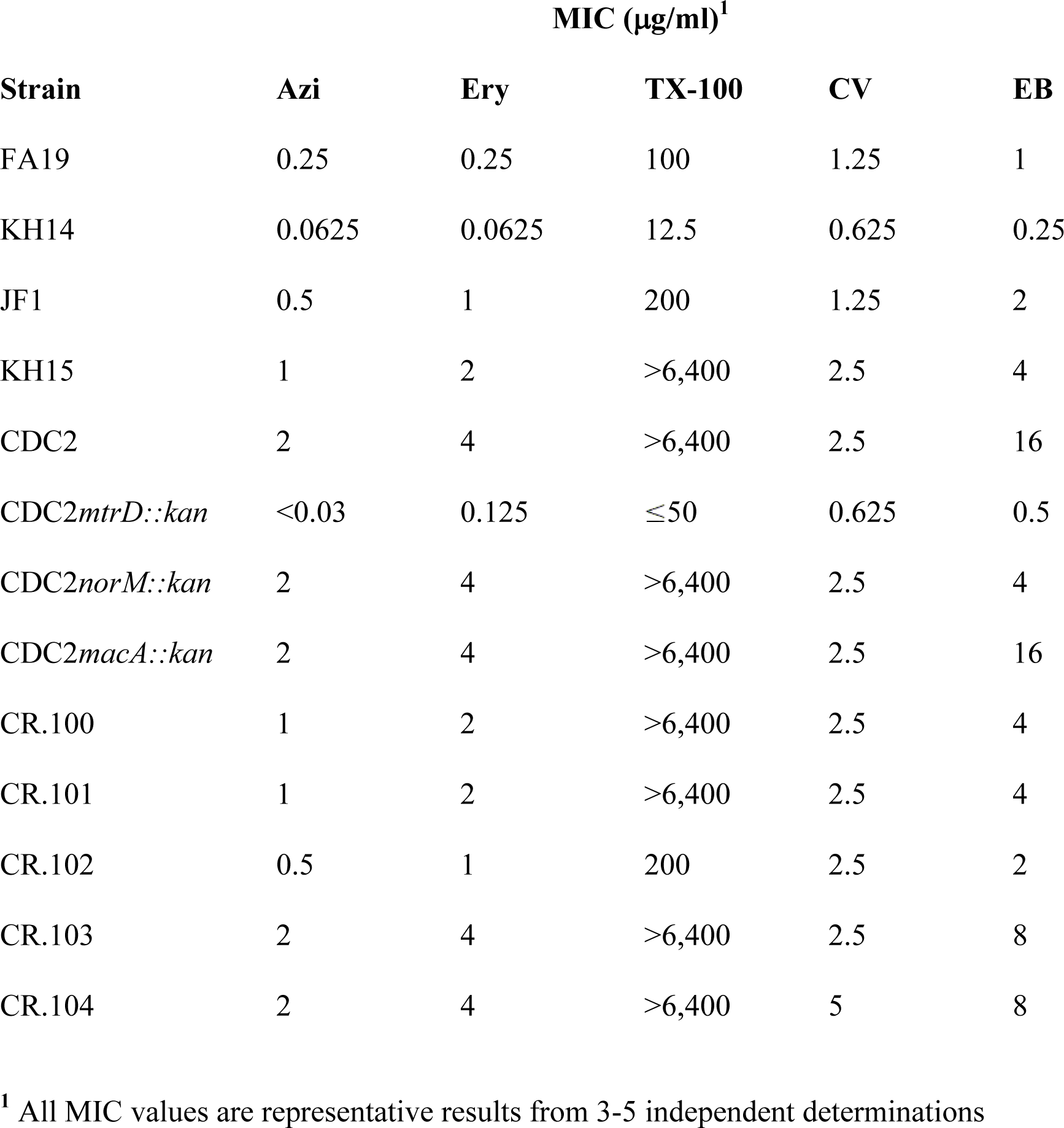
Levels of Antimicrobial Susceptibility of Gonococcal Strains

Since the MtrCDE efflux pump was essential for cross-resistance of CDC2 to antimicrobials, we focused on defining the impact of mosaic-like sequences in its *mtr* locus on MIC values of antimicrobials exported by this pump. For this purpose, we studied expression of the *mtrR* repressor gene and *mtrE* gene, which encodes the outer membrane protein (OMP) channel of the pump and is the last gene in the *mtrCDE* operon (Fig. 1 and ref. 3). Expression of these genes in FA19 and CDC2 was assessed at the levels of transcription and translation by qRT-PCR and western immunoblotting, respectively. At the level of transcription, it was found that *mtrR* expression in FA19 was significantly higher than that of CDC2, while *mtrE* expression was higher in CDC2 (Figure 2A). With respect to the MtrR repressor protein, its level in CDC2 and the other seven clinical Azi-alert strains was substantially less than that of strain FA19 (Figure S2A). As expected by the low level of the MtrR repressor in CDC2, the level of the MtrE OMP in this strain was higher than that of FA19. In this respect, the level of MtrE in CDC2 was similar to that of strain JF1 (FA19Δ*mtrR*), but less than that of KH15 (FA19 with single bp deletion in the *mtrR* promoter, which is known to decrease *mtrR* expression and increase *mtrCDE* expression (Fig. S2B).

**Figure 2.**
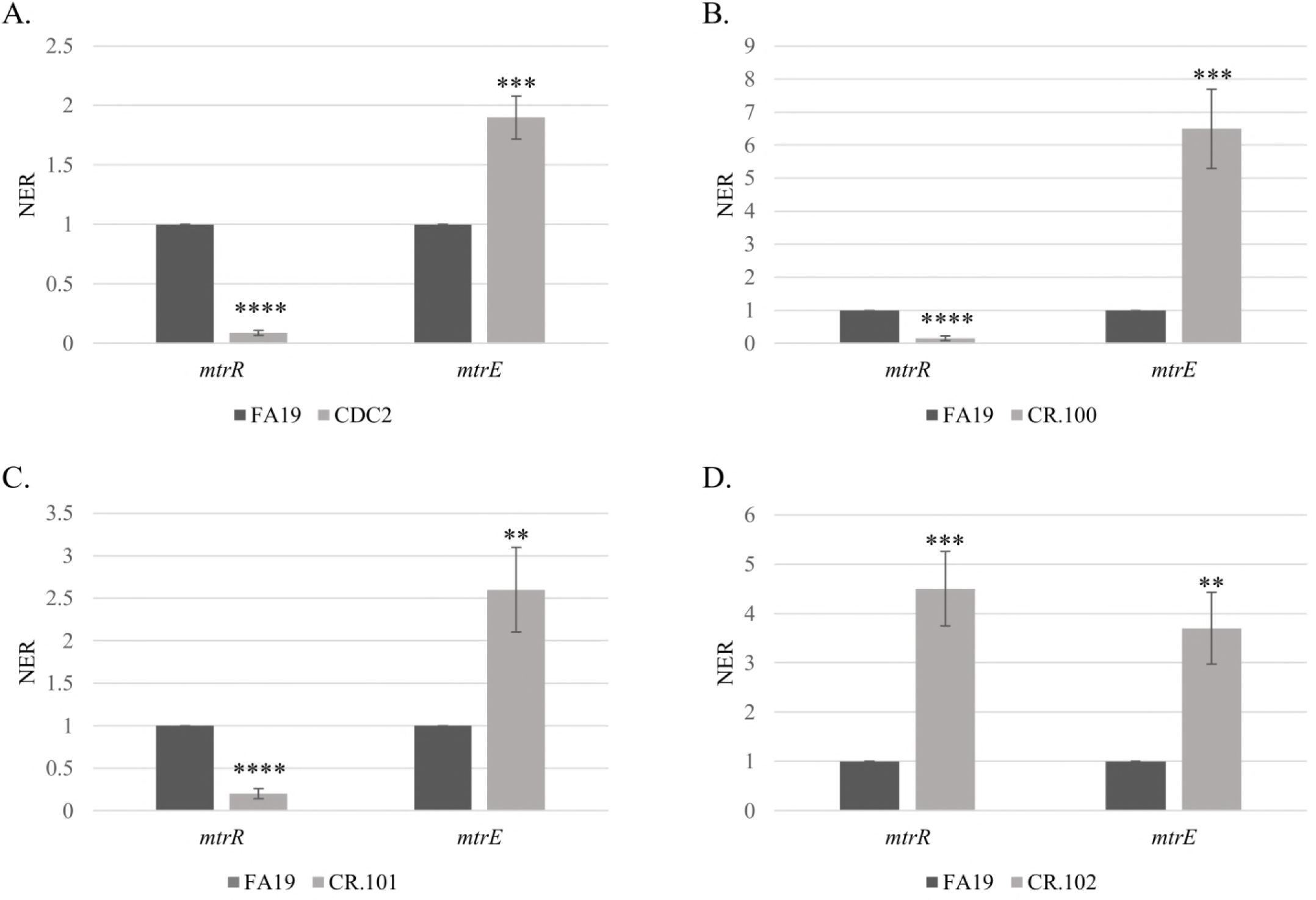
Shown are levels of expression of *mtrR* and *mtrE* genes in gonococcal strains FA19 and CDC2 (A), FA19 and CR.100 (B), FA19 and CR.101 (C) and FA19 and CR.102 (D). Gene transcript levels were quantified by qRT-PCR performed in triplicate with three biological replicates. Results are presented as average NER (normalized expression ratio) values (±SD) with P values. ** = P ≤0.01; *** = P ≤0.001; **** = P ≤0.0001

### Increased expression of *mtrCDE* due to mosaic-like *mtrR* and *mtrR-mtrCDE* promoter regions can contribute to, but is not sufficient, for the decreased Azi susceptibility phenotype of CDC2

We hypothesized that the less Azi-susceptible phenotype of CDC2 was due, in part, to enhanced levels of the MtrCDE efflux pump resulting from *cis*- and/or *trans*-acting mutations generated by the mosaic-like sequence that influence *mtrR* and *mtrCDE* expression. To test this hypothesis, we replaced the FA19 *mtrR* coding and *mtrR-mtrCDE* promoter sequences with that possessed by CDC2 and tested if this would influence expression of these genes as well as increasing MICs of Azi and other antimicrobials recognized by the MtrCDE efflux pump. For this purpose, a PCR-generated DNA fragment containing the *mtrR* coding region, the *mtrR-mtrCDE* intervening region and the 5′ -end of *mtrC* present in CDC2 (summarized in Fig. 3A) was used to transform strain FA19 for increased resistance to a known MtrCDE substrate, triton X-100 (TX-100). A resulting transformant termed CR.100 (Tables 1 and 2) was selected for more detailed studies. DNA sequencing revealed that CR.100 had *mtrR* coding and *mtrR*-*mtrCDE* promoter sequences that contained most, but not all, of the CDC2 donor mosaic-like DNA in this region (Fig. 3A). With respect to the nucleotide changes in the non-coding sequence, CR.100 had the CDC2 mosaic-like sequence that included one nucleotide change within the - 35 hexamer of the *mtrR* promoter (C to T) and one change within the -35 hexamer of the *mtrCDE* promoter (T to G) (Fig. 3A and Table 2) that have been observed in other strains (26, 30). In this respect, Wadsworth *et al* found that the -35 *mtrR* promoter mutation was (present in GCGS0402, while the -35 *mtrCDE* promoter mutation was present in all three of the *mtr* mosaic-like strains used in their study (30). Although a nucleotide difference was also noted in the previously identified transcriptional start site (TSS) for *mtrCDE* transcription (18), primer extension analysis revealed that transcription of *mtrCDE* in strains CDC2 and FA19 was similarly initiated (Fig. S3). CR.100 also contained the CDC2-derived mosaic-like sequence in *mtrR*, which was characterized by missense mutations in codons 79 (D79N), 183 (S183N) and 197 (M197I) (Fig. 3B).

**Figure 3.**
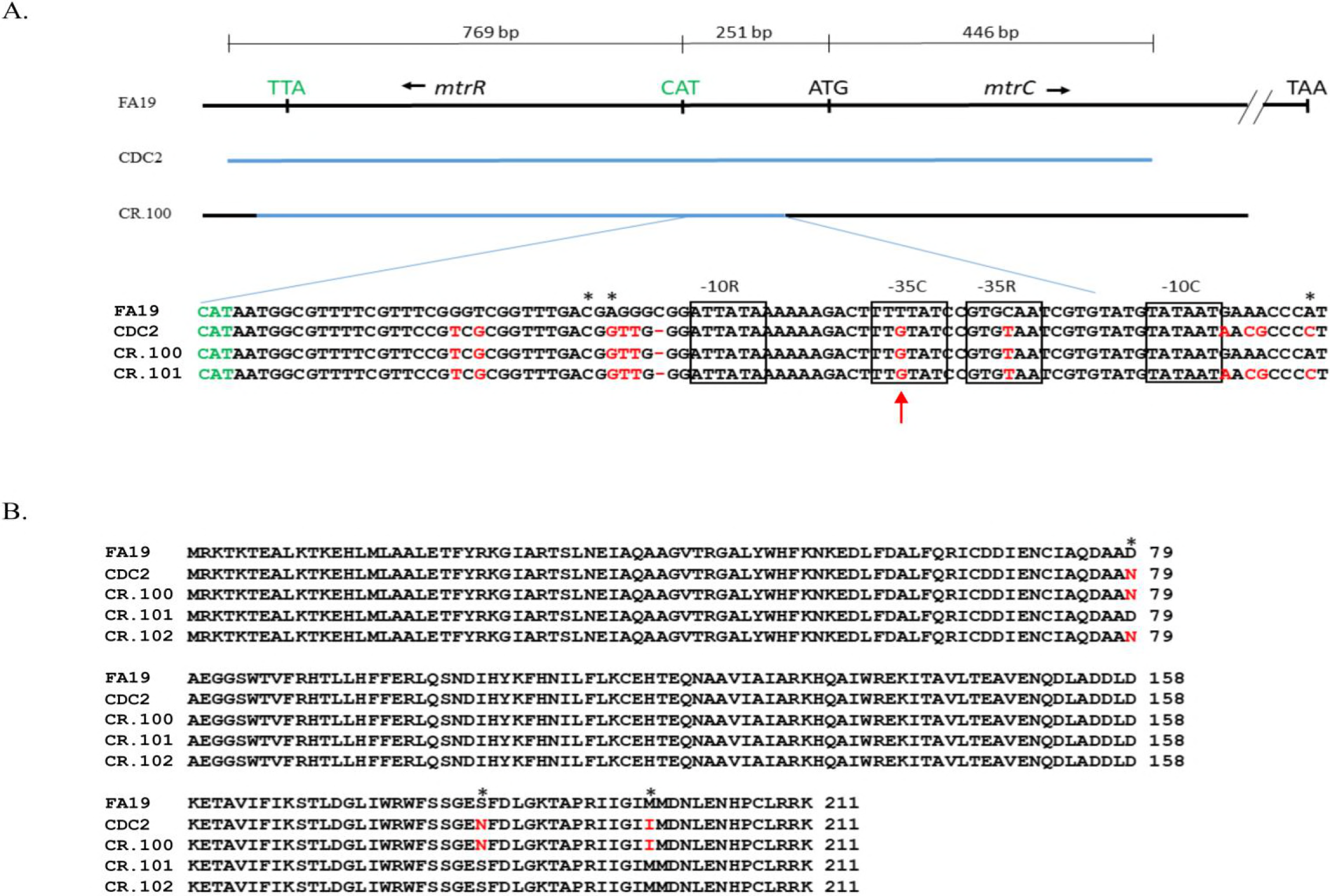
**(A)** Shown is the region of the *mtrR-mtrC* that was PCR-amplified from chromosomal DNA of strain CDC2 (blue line) used to transform strain FA19. The region of recombination in transformant strain CR.100 is shown by the blue line. The nucleotide sequences of the *mtrR*/ *mtrC* promoter region (*mtrCDE* coding strand) from strains FA19, CDC2, CR.100 and CR.101 are shown below. The translation start codon for *mtrR* is shown in green. The -10 and -35 hexamers of the *mtrR* and *mtrCDE* promoters are boxed. The TSS sites for both promoters are shown by asterisks. The red arrow shows the point mutation in the -35 hexamer of the *mtrCDE* sigma-70 promoter. Differences in nucleotide sequence or deletions are highlighted in red. (B) The predicted amino acid sequences of MtrR produced by strains FA19, CDC2, CR.100, CR.101 and CR.102 are shown. Differences at sites 79, 183 and 197 are highlighted in red and with asterisks.

**Table 2.** Summary of important *mtr* genetic changes and Azi and TX-100 MIC values

| Strain | MtrR | *mtrCDE* promoter/<br>*mtrR* promoter | MtrD | Azi | TX-100 |
| --- | --- | --- | --- | --- | --- |
| CDC2 | D79N, S183N,<br>M197I | -35 (T to G)/<br>-35 (C to T) | mosaic-like | 2 | >6,400 |
| FA19 | WT | WT/WT | WT | 0.25 | 100 |
| CR.100<sup>1</sup> | D79N, S183N,<br>M197I | -35 (T to G)/<br>-35 (C to T) | WT | 1 | >6,400 |
| CR.101 | WT | -35 (T to G)/<br>-35 (C to T) | WT | 1 | >6,400 |
| CR.102 | D79N | WT/WT | WT | 0.5 | 200 |
| CR.103 | D79N, S183N,<br>M197I | -35 (T to G)/<br>-35 (C to T) | 3'-end<br>mosaic-like | 2 | >6,400 |
| CR.104 | D79N, S183N,<br>M197I | -35 (T to G)/<br>-35 (C to T) | S821A,<br>K823E | 2 | >6,400 |

<sup>1</sup> Strains CR.100-104 are all in the FA19 genetic background

As is shown in Table 1, transformant strain CR.100 was more resistant than parent strain FA19 to a panel of MtrCDE substrate antimicrobials, but was two-fold less resistant than donor strain CDC2 to macrolides Azi and Ery and 4-fold less resistant to ethidium bromide (EB). An examination of transcript levels of *mtrR* and *mtrE* in FA19 vs. CR.100 showed that *mtrR* expression was decreased in CR.100 while *mtrE* expression was increased (Figure 2B). Although the MtrR repressor protein was readily detected in whole cell lysates of FA19, it was much lower in transformant strain CR.100 (Fig. 4). This result indicated that acquisition of the mosaic-like sequence encompassing the *mtrR* coding and *mtrR-mtrCDE* promoter sequences resulted in transcriptional repression of *mtrR* and de-repression of *mtrCDE*. However, it was unclear as to whether these gene expression differences and increased MICs of antimicrobials were due to the *mtrR*-coding or promoter mutations present in CR.100.

**Figure 4.**
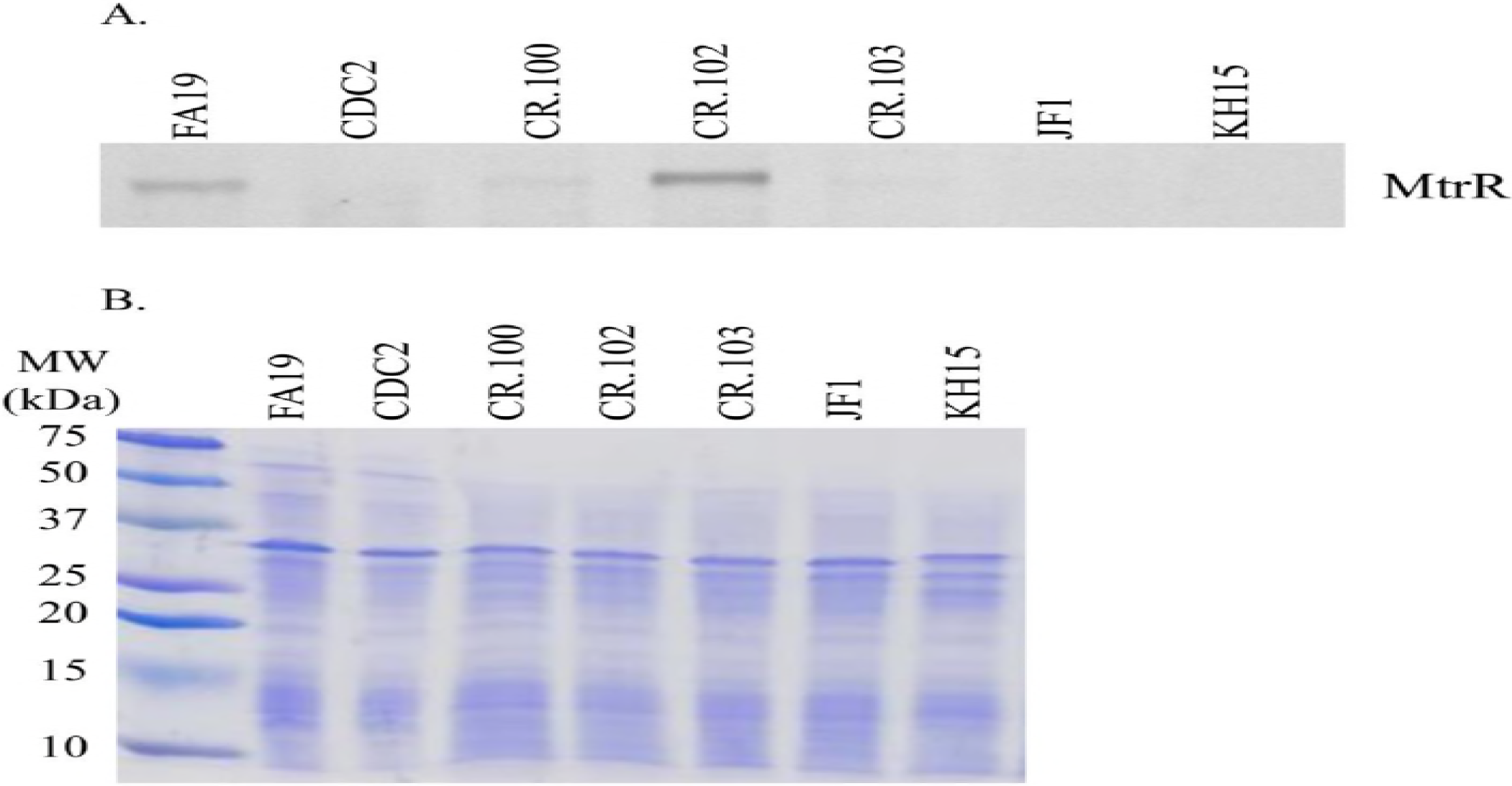
**(A)** Shown are levels of MtrR repressor protein in whole cell lysates of gonococcal strains as determined by western immunoblotting. (B) The SDS-PAGE gel stained with CBB showing near equivalent levels of protein (15 micrograms) loaded in each well is shown. Gonococcal strains are identified at the top of each well.

In order to separate potential influences of *cis*- or *trans*-acting mutations in the overlapping *mtrR*/*mtrCDE* promoter regions with that of the *mtrR* coding region, respectively, on levels of antimicrobial resistance and gene expression in CR.100, we generated a pair of PCR products that singularly covered these regions in CDC2. We found that both regions could transform WT strain FA19 for decreased susceptibility to TX-100 or Azi. Recovered transformants were termed CR.101 and CR.102. DNA sequencing of PCR products from CR.101 and CR.102 was performed to determine the extent of recombination of the donor mosaic-like sequence in the *mtrR* coding or upstream non-coding region. With the promoter region-bearing PCR product, we found that complete replacement of the wild-type *mtrR-mtrCDE* intervening region by the donor sequence from CR.100 had occurred in CR.101 (Table 2 and Fig. 3A). In contrast, with the *mtrR* coding PCR product, only the MtrR D79N mutation was present in CR.102; the MtrR amino acid alignment information is provided in Fig. 3B and summarized in Table 2. Antimicrobial susceptibility testing results (Table 1) showed that the MICs of macrolides against CR.101 (mosaic-like *mtrR-mtrCDE* intervening region) was two-fold higher than that of CR.102 (MtrR D79N). Interestingly, TX-100 resistance was > 32-fold higher in CR.101 than CR.102 (MtrR D79N); the MIC against CR.101 resembled that seen with KH15 (Table 1) which has a single bp deletion in the *mtrR* promoter that abrogates *mtrR* gene expression and shifts RNA polymerase recognition to the *mtrCDE* promoter (15). In contrast, the TX-100 MIC versus CR.102 was similar to that of JF1 (Table 1), which has a deletion of *mtrR* but retains a wild-type *mtrR* promoter (34).

The results from antimicrobial susceptibility testing suggested that *mtrR* coding and non-coding mosaic-like sequences have different impacts on expression of *mtr*-associated genes. Indeed, we found that while the MtrR D79N mutation in CR.102, which is also possessed by GCGS0402 and GCGS0834 (30), resulted in increased levels of *mtrE* expression compared to parental strain FA19 it also, unlike CDC2 and CR.100, had increased levels of the *mtrR* transcript (Fig. 2D) and the MtrR repressor protein (Fig. 4). Although position 79 of MtrR is outside of the DNA-binding domain of this repressor (14, 15), we suggest that this amino acid change causes a decrease in MtrR function resulting in de-repression of *mtrCDE* expression. Nevertheless, the consequence of this mutation did not endow gonococci with either the antimicrobial resistance profile or the *mtrR* gene expression profile observed for CDC2 or CR.100 that also have *mtrR*/*mtrCDE* promoter mutations (Table 1 and Fig. 2).

Based on the findings with CR.102, we next tested whether potentially *cis*-acting mutations in the *mtrR-mtrCDE* promoter region influence expression of *mtrR* or *mtrCDE*. As was observed with clinical isolate CDC2 and transformant strain CR.100, the presence of the mosaic-like *mtrR-mtrCDE* promoter region in CR.101 resulted in a decreased level of *mtrR* expression, but increased amounts of the *mtrE* transcript compared to parental strain FA19 (Fig. 2C). We hypothesize that mutations upstream of *mtrR*, especially the single nucleotide changes in the adjacent -35 hexamers of the *mtrR* and *mtrCDE* promoters (Fig. 3A), negatively impact expression of *mtrR* expression. Such repression of *mtrR* would not occur in CR.102 (MtrR D79N) where the wild-type FA19 promoter sequence is present (Fig. 3A) and MtrR levels are elevated (Fig. 4). Coupled with the antimicrobial susceptibility data (Tables 1 and 2), this result indicated that the nucleotide changes in the promoter region were responsible for modulating expression of *mtrR* and *mtrCDE* in clinical isolate CDC2 and transformant strain CR.100.

### A single nucleotide change in the mosaic-like *mtrR*/*mtrCDE* promoter region can impact gene expression

We hypothesized that the T to G change in the -35 hexamer region of the *mtrCDE* sigma-70 promoter in clinical isolate CDC2 and FA19 transformant strains CR.100 and CR.101 (red arrow, Figure 3A) directly raised *mtrCDE* expression (Figure 2A) and the MICs of MtrCDE antimicrobial substrates (Table 1); a similar nucleotide change was observed by others in *mtr* mosaic-like strains (26, 30). This nucleotide change would result in an improved -35 hexamer (5′-TT<u>T</u>TAT-3′ to 5′-TT<u>G</u>TAT-3′) of the *mtrCDE* promoter. To test the importance of this T to G change, two pLES94 *mtrCpromoter-lacZ* fusion (P_*mtrC-lacZ*_) constructs containing either the CDC2 *mtrCDE* promoter (CR.102pLES2.2) or an identical sequence but with the T nucleotide (CR.102pLES4.1) were introduced into CR.102 (FA19 MtrR D79N) by transformation; the *lacZ* fusions integrated in the *proAB* region of the gonococcal chromosome (35). After verification of selected transformants by DNA sequencing of PCR products, levels of beta-galactosidase (β-gal) were measured. As is shown in Fig. 5, the β-gal expression level was 3.5-fold higher in gonococci with the P_*mtrC-lacZ*_ fusion that contained the G nucleotide in the -35 sequence of *mtrCDE* promoter compared to the variant that had the T nucleotide. This result suggests that the T to G change observed in the -35 hexamer of the *mtrCDE* promoter possessed by CDC2 as well as other isolates (30) results in a more effective *mtrCDE* promoter than that possessed non-mosaic strain FA19. However, since MtrR can activate certain gonococcal genes (34) it was necessary to eliminate the (remote) possibility that the T to G nucleotide change facilitated binding of MtrR D79N to the *mtrCDE* promoter in an activating capacity. Accordingly, we introduced the P_*mtrC-lacZ*_ fusion in pLES2.2 into strain JF1 (FA19 Δ*mtrR*). The results showed that compared to CR.102, expression of P_*mtrC-lacZ*_ from the pLES2.2 fusion in JF1 was slightly elevated (data not presented), which is consistent with the MtrR D79N protein in CR.102 retaining a degree of transcriptional repressive activity as opposed to becoming an activator of *mtrCDE*.

**Figure 5.**
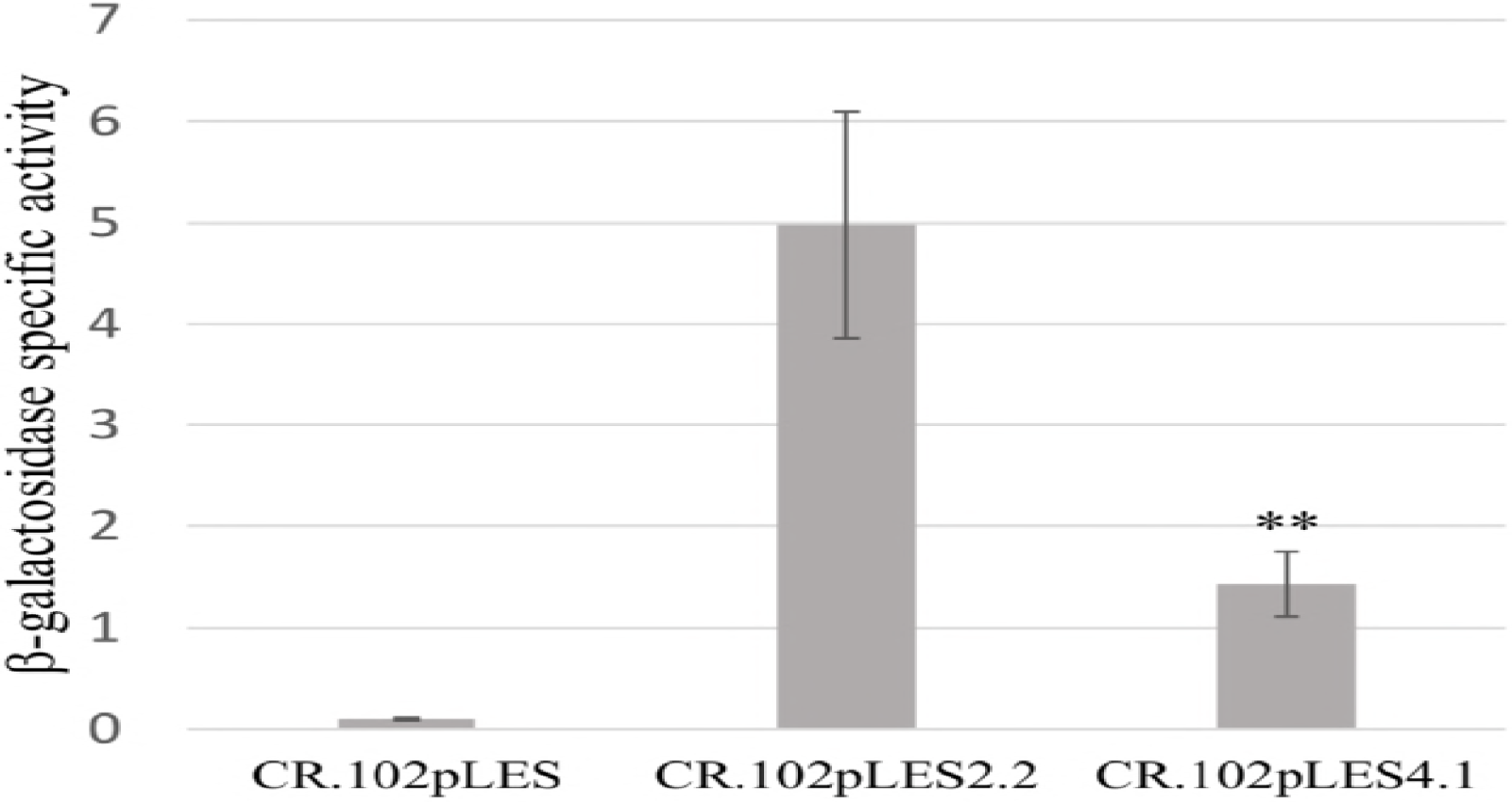
Shown are specific activities of β-gal produced by gonococcal strain CR.102 (MtrR D79N) from pLES94 constructs without P_*mtrC-lacZ*_ (CR.102pLES; control), or with P_*mtrC-lacZ*_with the CDC2 promoter (CR.102pLES2.2) or the same but with the WT -35 hexamer possessed by FA19 (CR.102pLES4.1). The results are shown as average values (±SD) with P values from three biologic replicates with each performed in triplicate. ** = P ≤0.01

### Mutations in *mtrD* due to acquisition of a mosaic-like sequence can increase antimicrobial resistance

The results from the gene expression studies described above implicated the T to G change in the - 35 *mtrCDE* promoter hexamer as having a strong cis-*acting* influence that de-represses *mtrCDE* expression leading to decreased susceptibility of gonococci to antimicrobials, including Azi. Collectively, these findings likely explain the levels of *mtrR* and *mtrCDE* gene expression in *mtr* mosaic-like strain CDC2 and other clinical isolates (26, 30) that possess this mutation. However, these transcriptional changes could not fully explain the higher MIC values of certain antimicrobials against CDC2. Thus, although transformant strain CR.100 had decreased susceptibility to all tested antimicrobials, only the MIC values of crystal violet (CV) and TX-100 matched that of CDC2 (Table 1). We hypothesized that amino acid differences in MtrD efflux pump transporter protein possessed by CDC2 compared to FA19 and CR.100 might account for the higher Azi MIC seen with strain CDC2. Based on the structure of MtrD (PDB 4MTI, ref. 36) and known similarity to AcrB (43), the following amino acid changes in MtrD possessed by strain CDC2 are predicted (E. Yu, personal communication) to impact its binding of antimicrobials: T42N, H46R, I48T, N101D, V662I, and K823E. To test whether these or other amino acid changes contribute to antimicrobial resistance in CDC2, three overlapping PCR products (summarized in Fig. S4) encompassing its entire *mtrD* gene were used together to transform strain CR.100 for resistance to 1 µg/ml of Azi. A resulting transformant (CR.103) was examined in antimicrobial susceptibility assays and was found to have identical levels of resistance as CDC2 to Azi (MIC of 2 µg/ml) and Ery (MIC of 4 µg/ml,), but was two-fold less resistant than CDC2 to the dye EB (Table 1). DNA sequencing of *mtrD* PCR products from CDC2, CR.100 and CR.103 showed that the nucleotide sequence of *mtrD* in these strains was identical to that of FA19 until codon 738 after which the CDC2 sequence had recombined in CR.103 until codon 1020. This recombination event generated twenty-three amino acid replacements in the C-terminal end of MtrD (Fig. 6). The amino-acid changes in this C-terminal region of MtrD are located in the DC sub-domain of the docking domain as well as in the PC2 sub-domain of the pore region and transmembrane domains TM8 and TM9 (36). As assessed by qRT-PCR, CR.100 and CR.103 did not differ in levels of the *mtrD* transcript (data not presented), indicating that differences in levels of antimicrobial resistance between these strains was linked to structural alterations of MtrD located at the C-terminal end.

**Figure 6.**
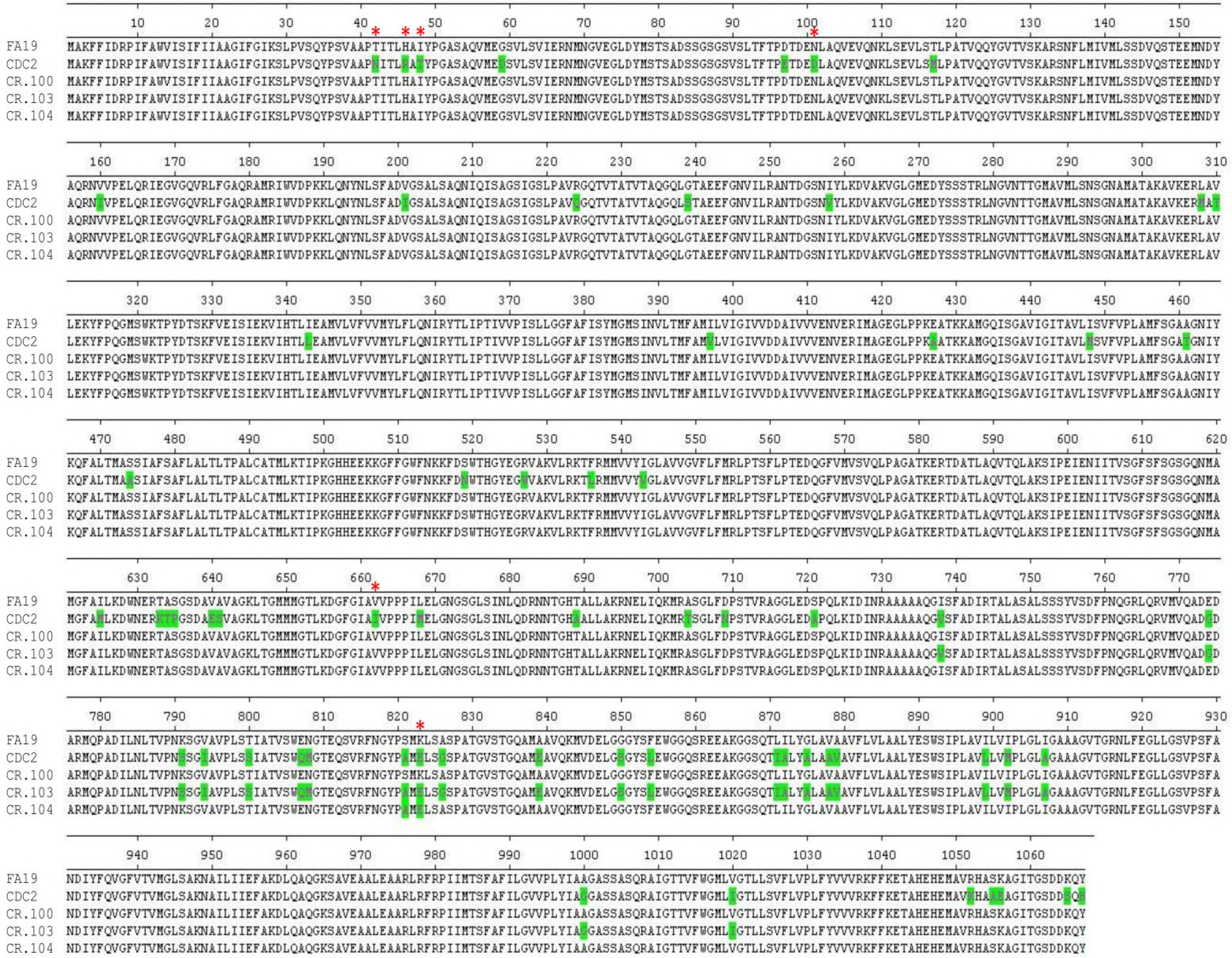
Shown are the sequences of the MtrD protein produced by gonococcal strains as deduced by DNA sequencing. Amino acid differences of MtrD from strains CDC2, CR.100, CR.103 and CR.104 compared to the FA19 are shown in green, with amino acids predicted to be sites for binding antimicrobials shown by red asterisks.

In order to verify that amino acid changes in the C-terminal region of MtrD from CR.103 could increase MIC values of Azi, we prepared an *mtrD* 3′-end PCR product described above from CR.103 and used it to transform CR.100 with selection for resistance to 1 µg/ml of Azi. Six individual transformants were assessed in antimicrobial susceptibility testing assays using Azi and Ery and all showed an elevated level of resistance to these macrolides comparable to that observed with donor strain CR.103 (data not presented). Differences were detected in the nucleotide sequence of the donated 3′-end of *mtrD* in the transformants indicating that unique sites of recombination had occurred. Translation of the nucleotide sequence to amino acid sequence showed similar as well as unique amino acid changes in each transformant strain (data not presented). Importantly, however, only two amino acid changes, positioned at 821 (Ser to Ala) and 823 (Lys to Glu), were common in all six transformants. In fact, one transformant (CR.104) had only these two amino acid changes (Fig. 6 and Table 2) compared to recipient strain CR.100. We also sequenced the entire *mtr* locus (6,793 bp) possessed by strains FA19 and CR.104 and found that except for the nucleotide differences in the *mtrR* coding, *mtrR*-*mtrCDE* intervening region and the *mtrD* allele of CR.104, the *mtr* locus in these strains were otherwise identical (data not presented). The MIC values of antimicrobials against CR.104 were identical to that of donor strain CR.103 (Table 1); the sole exception was with CV for which CR.104 was 2-fold more resistant.

## DISCUSSION

Recombination of commensal neisserial DNA sequences into the chromosome of *N*. *gonorrhoeae* and *N. meningitidis* has been well established (26, 30, 37-39). Such horizontal transmission of DNA can endow gonococci or meningococci with decreased susceptibility to antibiotics that target the respective gene product. The best example of the consequence of this genetic exchange event is that of mosaic sequences in the *pbp2* gene (also known as *penA*), which encodes the beta-lactam sensitive target, penicillin-binding protein 2 (PBP2) (40). The extensive re-modeling of PBP2 that occurs due to the multiple (up to 60) amino acid changes due to a mosaic *pbp2* decreases the acylation rate of penicillin and third generation cephalosporins (cefixime and ceftriaxone) (41). *pbp2* mosaic gonococcal strains also frequently contain *cis*-acting regulatory mutations that increase expression of the *mtrCDE* efflux pump operon (reviewed in [3]). Loss of the MtrCDE efflux pump in chromosomally-mediated penicillin resistant gonococci can result in a return to a penicillin-sensitive phenotype (29, 42), while a similar loss can result in a 2-4- fold increase in susceptibility to Cro and Azi (29). Thus, mutations that de-repress *mtrCDE* expression can work with mosaic *pbp2* to decrease gonococcal susceptibility to beta-lactams.

The mosaic-like *mtrR/mtrCDE* promoter region described herein and that reported by Wadsworth *et al.* (30) emphasizes that in gonococci nucleotide changes due to acquisition of mosaic-like *mtr* sequences can elevate *mtrCDE* expression and MICs of antibiotics; a graphic model describing the influence of these nucleotide changes was recently presented in a commentary (44) by the corresponding author (W.M.S.) on the Wadsworth et al. manuscript. We propose that the single nucleotide change (T to G) within the -35 hexamer of the *mtrCDE* promoter provides increased expression of *mtrCDE* as this change results in a consensus -35 hexamer sequence for sigma-70 promoters. Further, since this nucleotide change is also within the 13 bp inverted repeat positioned between the -10 and -35 regions of the *mtrR* promoter it could alter the ability of this sequence to form a DNA secondary structure with regulatory activity. Thus, the position of this nucleotide change would maintain the spacing between the -10 and -35 hexamers of the *mtrR* promoter but would reduce the T:A bp repeat from 6 to 4. This reduction in the T:A bp repeat may impact promoter recognition by RNA polymerase, which preferentially shifts its recognition from the *mtrR* promoter to the *mtrCDE* promoter that has an improved -35 hexamer sequence. It should also be noted that *mtr* mosaic-like CDC2 exhibited a low level of *mtrR* expression possibly due to the nucleotide change in the –35 hexamer of the *mtrR* promoter (Figs. 2 and 3). We hypothesize that this may also contribute to the low levels of MtrR in this and other clinical isolates. Interestingly, when the MtrR D79N mutation from CDC2 was placed into strain FA19, which has a WT *mtrR* promoter, levels of MtrR increased significantly (Fig. 4) indicating that this mutation can cause dysregulation of autoregulation of *mtrR* expression.

While the -35 *mtrCDE* promoter mutation can endow gonococci with increased expression of *mtrCDE* (Fig. 2) and elevate gonococcal resistance to antimicrobials, it did not account for the Azi MIC observed with clinical isolate CDC2 (Tables 1 and 2). In this respect, the results presented herein and that of Wadsworth *et al.* (30) show that amino acid changes in *mtrD,* which result from importation of commensal neisserial DNA, are also necessary for the less Azi-susceptible phenotype of *mtr* mosaic-like strains such as CDC2 and GCGS0402. Based on the published MtrD structure (36), many of the amino acid positions in MtrD that are changed in CDC2 versus antibiotic-sensitive strain FA19 could influence drug binding, efflux activity or protein stability. Results from our transformation experiments indicated, however, that amino acid changes in the C-terminal end of MtrD, especially at positions 821 (S821A) and 823 (K823E), are sites for gain-of-function mutations that can contribute to antimicrobial resistance seen in *mtr* mosaic-like strains. These two amino acids are within the PC2 region of MtrD that is part of the pore domain of this transporter and is located at the outermost region that faces the periplasm. Based on the structural comparison of MtrD with the orthologous AcrB protein (44), position 823 in MtrD is predicted to be a site for binding antimicrobials (E. Yu, personal communication). Amino acid changes located elsewhere in MtrD could also have a similar impact, as indicated by the higher EB MIC value against CDC2 versus transformant strains CR.103 and CR.104. We note that the S821A and K823E changes are also present in the MtrD protein of the GCGS0402 and GCGS0834 strains studied by Wadsworth *et al.* (30); interestingly, GCGS0276, that also possesses a mosaic-like *mtrD,* has a K823D change that may have a similar functional impact on MtrD activity as the K823E mutation (30).

Active international and national surveillance systems for determining trends in gonococcal resistance to antibiotics coupled with whole genome sequencing and bioinformatic analyses have been instrumental in detecting gonococci with resistance determinants including those generated by mosaic-like *mtr* sequences (45-49). The detailed molecular, genetic and phenotypic analyses made possible by these efforts have provided new insights regarding the impact of the gonococcal MtrCDE efflux pump in determining levels of bacterial resistance to clinically important antibiotics. The world-wide distribution of strains with mosaic-like *mtr* loci indicates that future diagnostic tests for resistance determinants should include functionally important nucleotide changes that were not described in earlier studies (summarized in [13]). Further, the presence of such gonococcal strains re-emphasizes that genetic exchange between gonococci and commensal *Neisseria*, which is likely to occur in the pharyngeal cavity, can be a major mechanism by which *N. gonorrhoeae* develops resistance to antibiotics. Thus, in order to enhance monitoring of antibiotic resistance trends, it would be prudent to routinely sample both genital and extra-genital (especially the oral mucosae) sites for the presence of gonococci in patients suspected of having gonorrhea so as to better detect emergence of antibiotic resistant strains in the community.

## MATERIALS AND METHODS

### Gonococcal strains, growth conditions, oligonucleotide primers and determination of susceptibility to antimicrobial agents

The gonococcal strains used in this study are presented in Tables S1 and S2. The clinical strains used in this investigation were kindly provided by Edward Bannister, PhD (Dallas, TX) and the Dallas, TX GISP sentinel site. These strains were provided to CDC without patient identifiers and as such, there was no involvement of human subjects in the research reported herein. For most genetic studies we used antimicrobial-sensitive strain FA19 with an introduced *rpsL* mutation (FA19 Str^R^) that confers high level resistance to streptomycin, but for brevity it is referred to as FA19 throughout the text (Table S2). Eight gonococcal clinical strains displaying decreased Azi-susceptibility between 2014 and 2015 were also included in this study (Table S1). The details of their WGSs and bioinformatic analysis will be presented elsewhere (Soge and McLean, in preparation). Briefly, the eight strains were selected based on their representation within one of four different clades that were identified by analysis of a generated phylogentic tree from thirty-seven strains. Of these, strain LRRBGS0002 (CDC2; Table S2) was the most extensively studied. Gonococcal strains were grown overnight at 37°C under 5 % (v/v) CO_2_ on GCB agar containing defined supplements I and II (21). The sequences of oligonucleotide primers used in this study are shown in Table S2. The MICs of antimicrobials were determined by the agar dilution method (50) using the GISP protocol (51). Antimicrobials were purchased from Sigma Chemical Co. (St. Louis, MO).

### Bioinformatic analyses of WGS information

To compare the sequences of the *mtr* loci among the strains, sequences for strains FA19, FA1090, and MS11 were retrieved from NCBI genome sequence database (https://www.ncbi.nlm.nih.gov/nuccore/) or NCBI Sequence Read Archive (https://www.ncbi.nlm.nih.gov/sra); the H041 WGS sequence was kindly provided by R. Nicholas (University of North Carolina-Chapel Hill, NC). Sequence read files for LRRB strains used in this study (Table S1) were downloaded from NCBI Sequence Read Archive (https://www.ncbi.nlm.nih.gov/sra) into CLC Genomics Workbench (https://www.qiagenbioinformatics.com/), where we performed trimming, *de novo* assembly, and a BLAST of the FA19 *mtr* locus sequence (GenBank accession number CP012026.1, nucleotides 1104741-1111533) against the newly assembled sequences in order to recover their *mtr* locus sequences for alignment. Raw sequence files for GCGS0276, GCGS0402, and GCGS0834 (30) were downloaded from NCBI SRA, and trimmed and assembled according to published methods with cutadapt (52) and SPAdes (53). NCBI-blast toolkit (BlastN [https://www.blast.ncbi.nlm.nih.gov]) retrieved the sequences for the *mtr* operon based on FA19 *mtr* locus sequence. Alignments were performed using the Clustal Omega multiple sequence alignment tool for nucleotide and amino acid sequences (54, 55), and percent identity values were obtained from the resulting pairwise identity matrix. The Newick file derived from the alignment was visualized using Interactive Tree of Life (iTOL) (56).

### Strain constructions

To construct mutants of the eight decreased Azi-susceptible clinical isolates genomic DNA from strains KH14 (FA19*mtrD::kan*), FA19 *norM::kan* and FA19 *macA::kan* were used in transformation experiments as described previously (21) and verified by PCR using oligonucleotide primer pairs for each gene (Table S3). Strains KH14 and FA19 *norM::kan* have been previously described (21, 57). FA19*macA::kan* was constructed in this study. Briefly, primers macAF and macAR (Table S2) were used to PCR amplify *macA* from FA19 genomic DNA. The resulting PCR product was cloned into the pBAD-TOPO vector as described by manufacturer (Invitrogen) to create pBAD *macA,* which was then digested with SmaI. The non-polar kanamycin (Kan) resistance cassette *aphA3* (58) was cloned into the SmaI site and the resulting plasmid (pBAD*macA::kan)* was transformed into FA19. Transformants were selected on GCB agar containing 50 μg/ml of kan and were verified by PCR. PCR products were generated using CDC2 genomic DNA and primers CEL1 and KH9#12B (to construct CR.100) or primers CEL4 and mtrCpromR (to construct CR.101) or primers CEL1 and KH9#10B (to construct CR.102). The resulting PCR products were transformed into FA19 as previously described (18). CR.100 clones were selected on GC plates supplemented with 3,600 μg/ml of TX-100. CR.101 was selected on GC plates supplemented with 0.25 μg/ml of Azi. CR.102 was selected on GC plates supplemented with 400 μg/ml of TX-100. The presence of mutations in the clones was verified by sequencing. For construction of CR.103, PCR products were generated with primers mtrD11Rev and mtrD3, mtrD3Rev and mtrD1, and mtrE12 and mtrD10 using CDC2 genomic DNA as template. The 3 PCR products were used together to transform strain CR.100 using a selection of 1 μg/ml of Azi; a resulting transformant was termed CR.103. For construction of CR.104, a PCR product was generated with primers mtrD10 and mtrE12 on genomic DNA from CR.103. The resulting PCR was transformed into CR.100 and clones were selected on GC agar plates supplemented with 1 μg/ml of Azi.

### Quantitative reverse transcriptase-polymerase chain reactions (qRT-PCR)

RNA was extracted from strains FA19, CDC2, CR.100, CR.101, CR.102 and CR.104 at mid-logarithmic phase of growth in GC broth plus supplements by the TRIzol method as directed by the manufacturer (Thermo Fisher Scientific, Waltham, MA) and was performed as described (59). Briefly, genomic DNA (gDNA) was removed by RNAse-free DNAse treatment and gDNA Wipeout (Qiagen, Germantown, MD). The resulting RNA was then reverse transcribed to cDNA using the QuantiTect Reverse Transcriptase kit (Qiagen) as described (57). Primers 16Smai_qRTF and 16Smai_qRTR were used for 16S rRNA. Primers mtrEqPCR-F and mtrEqPCR-R were used for the *mtrE* gene, primers mtrD8 And mtrD13 were used for the *mtrD* gene. Primers mtrR_qRT_F and mtrR_qRT_R were used for the *mtrR* gene. Sequences of primers are shown on Table S2. Results were calculated as normalized expression ratios (NER) using 16S rRNA expression levels. Statistical significance was calculated by Student’s t-test.

### Western immunoblot analysis

Whole-cell lysates were prepared from gonococcal strains grown overnight on GC agar plates with supplements as described previously and separated by sodium dodecyl sulfate polyacrylamide gel electrophoresis (SDS-PAGE) (60). Coomassie brilliant blue (CBB) staining of duplicate SDS-PAGE gels was performed to calibrate and verify consistent loading of proteins (15 µgs) loaded into each well. The concentration of protein in whole cell lysates was estimated by using a Nanodrop spectrometer at 280 nM. Western immunoblotting using a mouse anti-MtrE serum (kindly provided by A. Jerse) and a rabbit anti-MtrR serum was performed as described previously (23, 34).

### β-gal assays

*lacZ* fusions were constructed using the pLES94 system as previously described (35). Briefly, PCRs were performed on genomic DNA from strain CDC2 using primers C2 and C3PmtrC for pLES2.2 and C4 and C3PmtrC for pLES4.1. The resulting PCR products were cloned into the BamH1 site of pLES94. pLES94, pLES2.2 and pLES4.1 and were introduced into strains CR.102 or JF1 by transformation with selection on GC agar plates supplemented with 1 µg/ml of chloramphenicol. β-gal assays were performed in triplicate as described by Folster *et al.* (34) from lysates of gonococcal strains after growth overnight on GC agar plates supplemented with chloramphenicol. The β-gal specific activity was calculated using the formula: A_420_ x 1000/A_280_ (mg/ml) x time (min) x volume (ml).

### Primer extension analysis

Total RNA from strains FA19 and CDC2 was prepared from mid and late-logarithmic phase GCB broth cultures by the TRIzol method as directed by the manufacturer (Thermo Fisher Scientific, Waltham, MA). Primer extension experiments were performed as described previously (18) on 6 µg of total RNA with primer PEmtrC181. Primer extension transcription start site of the *mtrC* gene was determined by electrophoresis of the extension products on a 6% (w/v) DNA sequencing acrylamide gel adjacent to reference sequencing reactions.

## ACKNOWLEDGEMENTS

We thank E. Yu (Case Western University) for his insightful comments regarding MtrD structure and impact of mutations with respect to binding antimicrobials. We would also like to thank the Gonococcal Isolate Surveillance Project (GISP) for the use of isolates and corresponding data for this analysis. Edward Bannister, PhD (Dallas, TX) and the Dallas, TX GISP sentinel site collected, isolated and provided epidemiological data for the isolates used in this analysis. The University of Washington GISP Regional Laboratory (Seattle, WA) determined and provided antimicrobial susceptibility result data for the isolates used in this analysis. We also thank Steven Johnson, Hsi Liu, Matthew Schmerer, Sandra Seby, Jesse Thomas, and Eshaw Vidyaprakash for thoughtful and informative discussions. The contents of this article are solely the responsibility of the authors and do not necessarily reflect the official views of the National Institutes of Health, the Centers for Disease Control and Prevention, the U.S. Department of Veterans Affairs, or the United States government. We have no competing interest to declare.

---

**Table S1.**
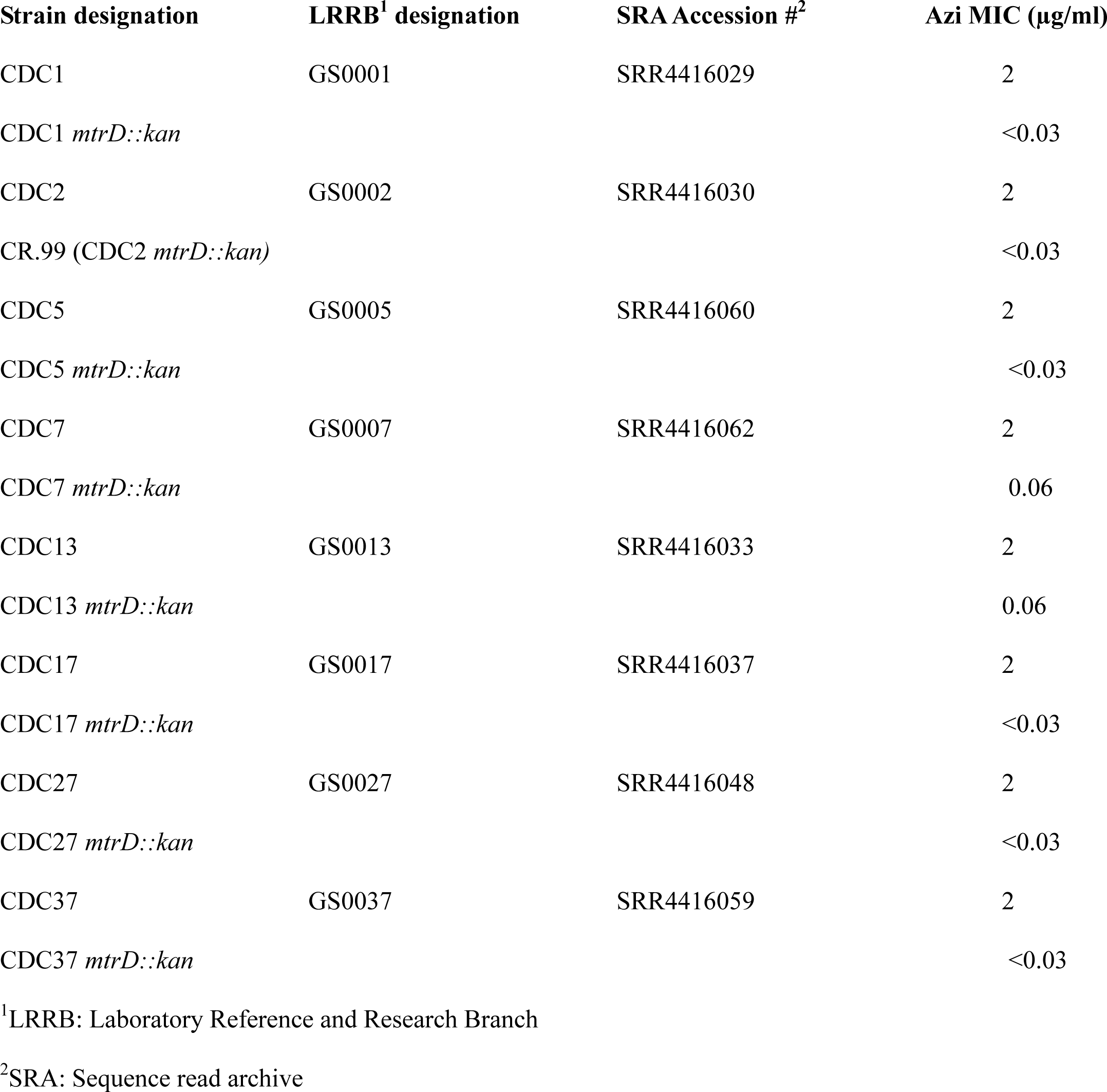
Gonococcal clinical strains and *mtrD::kan* mutants used in this study.

**Table S2.** Genetic derivatives used in this study.

| 751 | Strain | Genotype | Source |
| --- | --- | --- | --- |
| 752 | FA19Str<sup>R</sup> | point mutation in *rpsL* | Jerse, 2003 |
| 753 | JF1 | as FA19 with *mtrR* deleted | Hagman 1995 |
| 754 |  |  | Folster 2005 |
| 755 | KH14 | as FA19 but with *mtrD::kan* | Hagman 1997 |
| 756 | KH15 | as FA19 with 1 bp deletion in the |  |
| 757 |  | *mtrR* promoter | Hagman 1995 |
| 758 | CR.99 | as CDC2 but with *mtrD::kan* | this study |
| 759 | CR.100 | as FA19Str<sup>R</sup> with *mtrR* gene from |  |
| 760 |  | CDC2 and *mtrC* and *mtrR* promoters |  |
| 761 |  | from CDC2 | this study |
| 762 | CR.101 | as FA19Str<sup>R</sup> with *mtrR/mtrC* intergenic |  |
| 763 |  | region from CDC2 | this study |
| 764 | CR.102 | as FA19Str<sup>R</sup> with D79N mutation |  |
| 765 |  | in MtrR | this study |
| 766 | CR.103 | as CR.100 with *mtrD* mutations |  |
| 767 |  | (positions 2196 to 3072) from CDC2 | this study |
| 768 | CR.104 | as CR.100 with K823E mutation in MtrD | this study |
| 769 | FA19*norM::kan* | as FA19 with *norM::kan* | Rouquette- |
| 770 |  |  | Loughlin 2003 |
| 771 | FA19*macA::kan* | as FA19 with *macA::kan* | this study |
| 772 | CR102 pLES94, | control plasmid | this study |
| 773 | CR102 pLES2.2 | with *pmtrC-lacZ* CDC2 promoter | this study |
| 774 | CR.102 pLES4.1 | with *pmtrC-lacZ* CDC2 promoter with G to T change | this study |

**Table S3.**
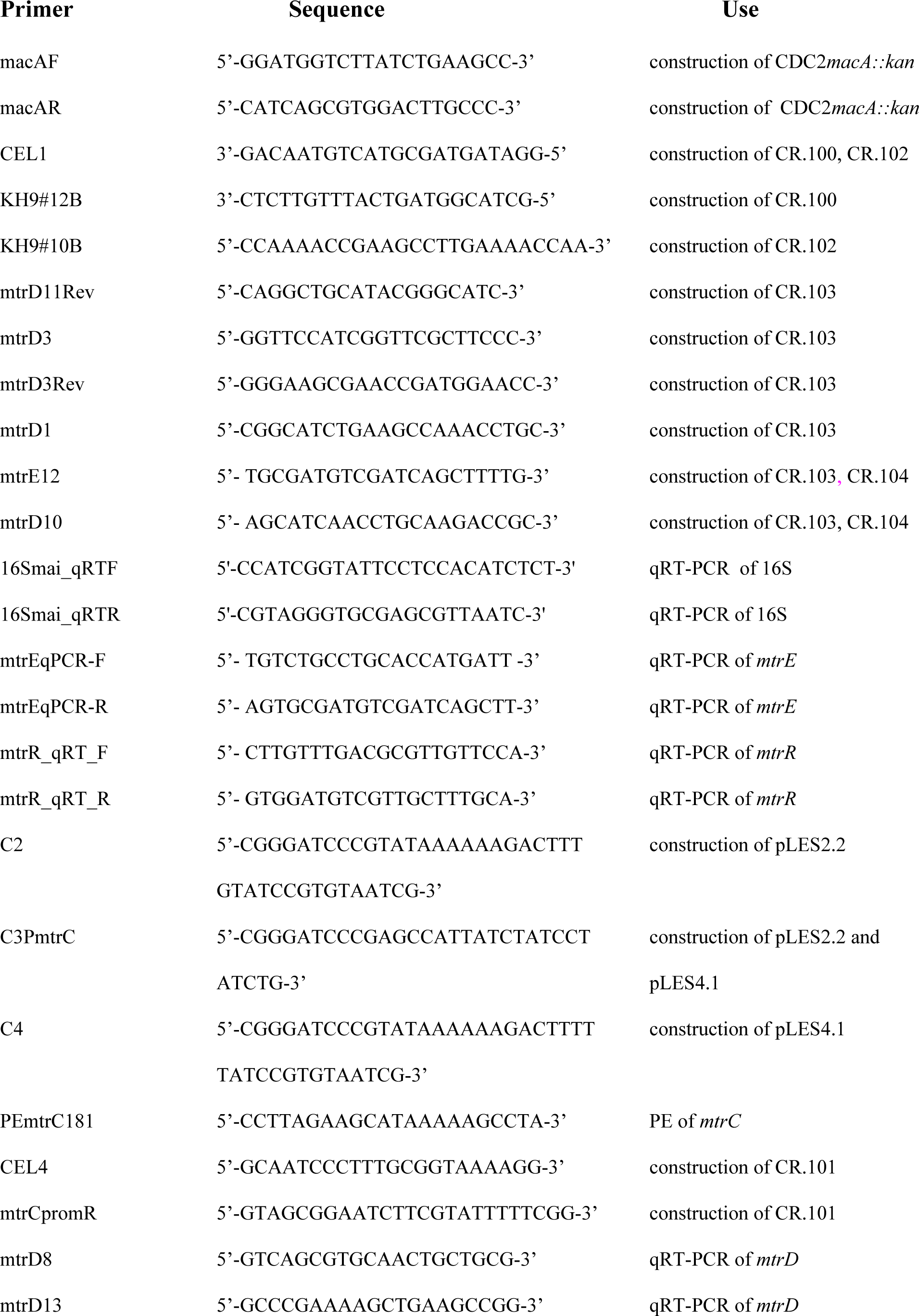
Sequences of oligonucleotide primers.

**Figure S1.** (A) Shown is an alignment of the nucleotide sequences of the complete *mtr* locus possessed by gonococcal strains FA19, CDC2 and three clinical isolates studied by Wadsworth *et al.*(30). Nucleotides of note in CDC2 that differ from FA19 in either coding or non-coding regions are highlighted in red. The Correia element (CE) in strain GCGS0276 is highlighted in gray. (B) Shown is a phylogenetic tree for the *mtr* loci possessed by gonococcal strains FA19, CDC2, GCGS0276, GCGS0402 and GCGS0834.

**Figure S2.**
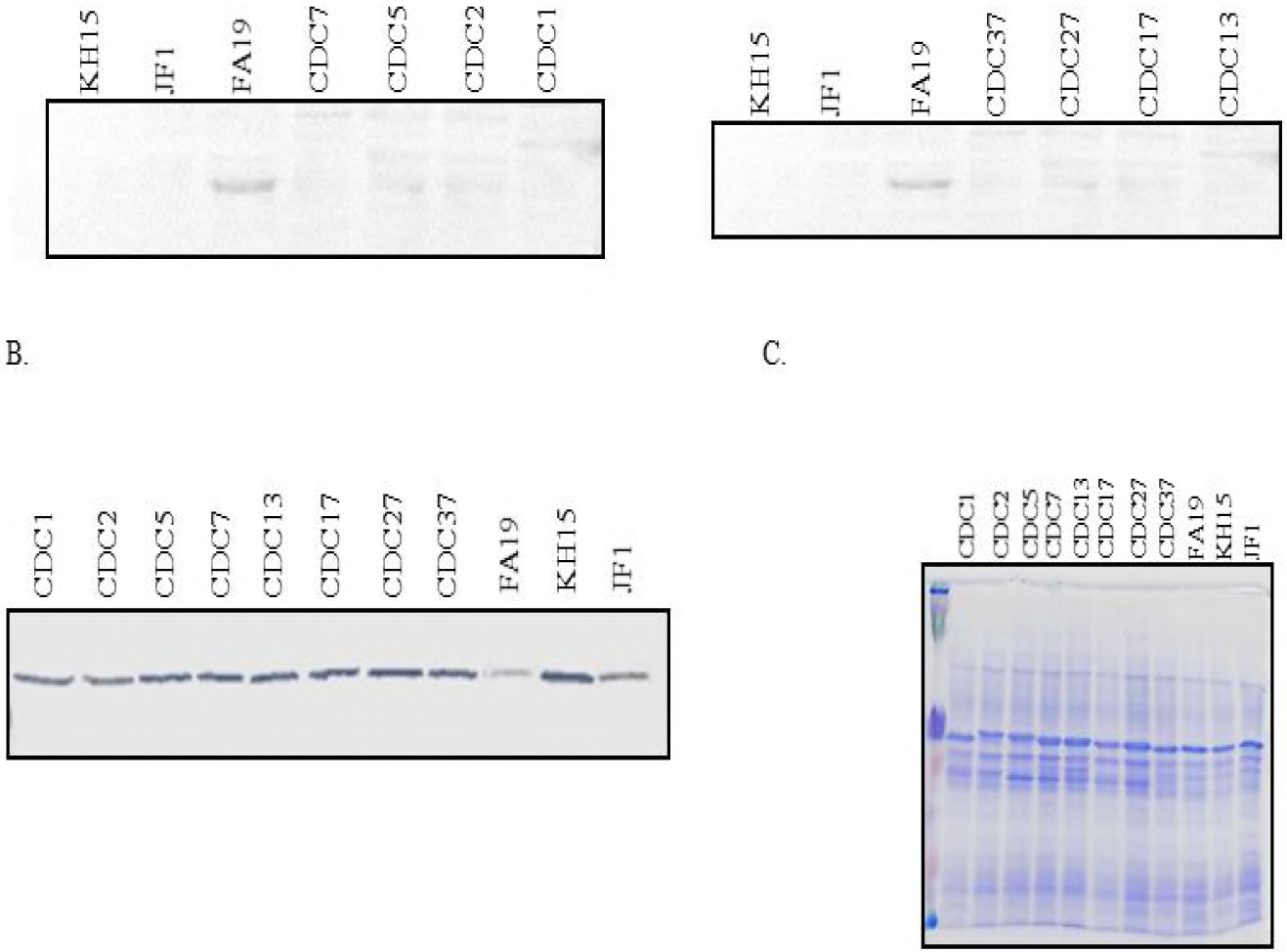
Shown are levels of the MtrR (A) and MtrE (B) proteins in whole cell lysates of gonococcal strains as determined by western immunoblotting. The eight CDC alert strains are shown with a strain number. Included in these blots are lysates from WT strain FA19 and transformant strains JF1 and KH15 that lack MtrR due to deletion of the gene (JF1) or a single bp deletion in the *mtrR* promoter that abrogates *mtrR* gene expression and elevates *mtrCDE* expression. An accompanying CBB-stained gel is shown panel C.

**Figure S3.**
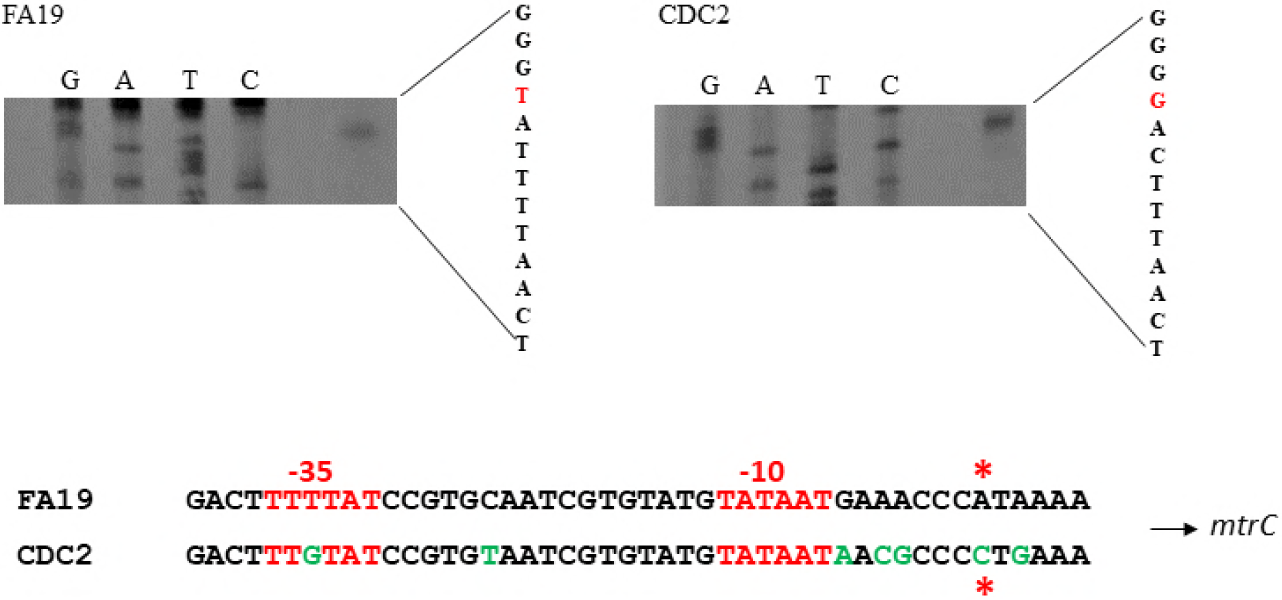
(A). Shown are results from a primer extension experiment that identified the *mtrC* TSS in gonococcal strains FA19 and CDC2. The nucleotide sequence from the noncoding strand is shown adjacent to autoradiogram with the start sites highlighted in red. (B). Shown are the nucleotide sequences of the *mtrCDE* promoter region from strains FA19 and CDC2 with the G nucleotide change (CDC2) in the -35 hexamer shown in green and the TSS sites highlighted by red asterisks.

**Figure S4.**
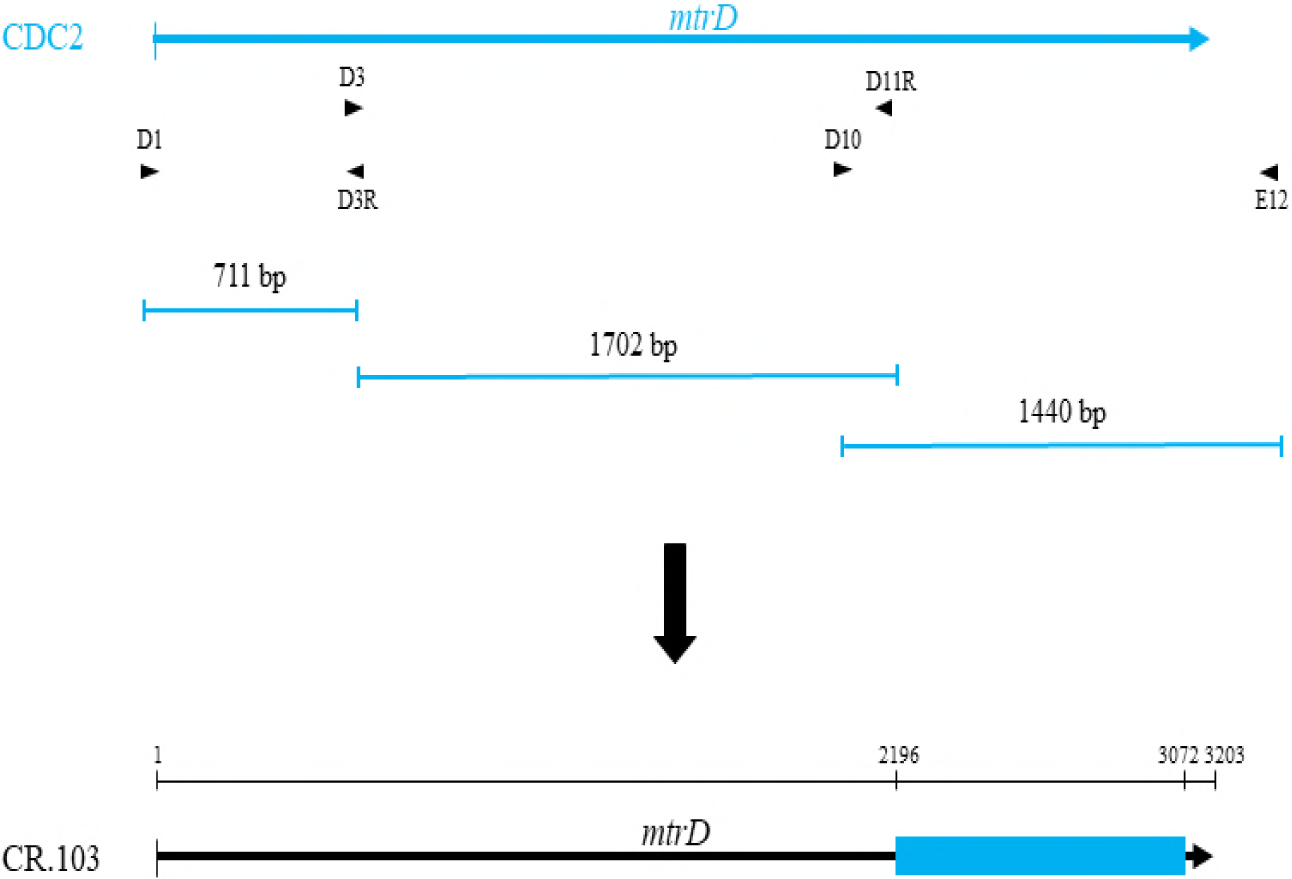
Shown is the strategy used to construct CR.103. Three regions of *mtrD* from CDC2 were amplified by PCR. The oligonucleotide primers and the length of the products are shown. These PCR products were used to transform strain CR.100 for resistance to 1 µg/ml of Azi. The region of recombination in strain CR.103 is shown by the blue rectangle.

